# A dorsal versus ventral network for understanding others in the developing brain

**DOI:** 10.1101/2024.05.23.595476

**Authors:** Clara Schüler, Philipp Berger, Charlotte Grosse Wiesmann

## Abstract

Young children strongly depend on others, and learning to understand their mental states (referred to as Theory of Mind, ToM) is a key challenge of early cognitive development. Traditionally, ToM is thought to emerge around the age of 4 years. Yet, in non-verbal tasks, preverbal infants already seem to consider others’ mental states when predicting their actions. These early non-verbal capacities, however, seem fragile and distinct from later-developing verbal ToM. So far, little is known about the nature of these early capacities and the neural networks supporting them. To identify these networks, we investigated the maturation of nerve fiber connections associated with children’s correct non-verbal action prediction and compared them with connections supporting verbal ToM reasoning in 3- to 4-year-old children, that is, before and after their breakthrough in verbal ToM. This revealed a ventral network for non-verbal action prediction versus a dorsal network for verbal ToM. Non-verbal capacities were associated with maturational indices in ventral fiber tracts connecting regions of the salience network, involved in bottom-up social attention processes. In contrast, verbal ToM performance correlated with maturational indices of the arcuate fascicle and cingulum, which dorsally connect regions of the default network, involved in higher-order social cognitive processes including ToM in adults. As non-verbal tasks were linked to connections of the salience network, young children may make use of salient perceptual social cues to predict others’ actions, questioning theories of mature ToM before 4 years.

**Significance:** As highly social beings, humans frequently reason about other people’s thoughts, termed Theory of Mind (ToM). While ToM is traditionally assumed not to emerge before 4 years, preverbal infants already seem to consider others’ thoughts when predicting their actions non-verbally. This raises the question of when ToM develops and what explains this discrepancy. We show that young children’s success in non-verbal tasks is related to different neural networks than those involved in mature verbal ToM. While verbal ToM was linked to ToM network connections, younger children’s non-verbal capacities were associated with the maturation of connections of the salience network. This indicates that, instead of mature ToM, young children might utilize salient social cues to predict others’ actions.

## Introduction

Humans are highly social beings, and in social contexts, frequently infer what other people think or believe, referred to as Theory of Mind (ToM) (1). In children’s cognitive development, the emergence of ToM thus marks a crucial milestone. The traditional view posits that ToM relies on higher cognitive function and, thus, only emerges around 4 years when children begin to verbally reason about others’ mental states (2). This view, however, seems inconsistent with non-verbal tasks showing that preverbal infants already consider others’ mental states when predicting others’ actions (including their intentions, perspectives, knowledge, and possibly even beliefs) (3, 4). Do infants already have a ToM, and do they reason about others’ mental states as adults and older children do? Recent studies cast doubt on this idea by showing that success in non-verbal ToM tasks is fragile (e.g., (5, 6), unrelated to later-developing verbal ToM (7, 8), and dissociated from classic ToM regions in the brain (9). This suggests that infants’ correct action prediction in non-verbal ToM tasks relies on a different system than mature verbal ToM, raising the question of which processes underlie these early capacities. Understanding the neural networks involved in young children’s correct non-verbal action prediction can shed light on the underlying processes. Here, we set out to identify the networks that support non-verbal action prediction in children before and after their breakthrough in verbal ToM reasoning (i.e., 3 to 4 years old) and compare these networks with those involved in verbal ToM in the same individuals.

In the traditional verbal ToM tasks, children are asked how an agent with a false belief will act, for example, where they will search for an object whose location they don’t know (2). In contrast, in the non-verbal tasks, children are not directly asked about the agent’s belief or actions. Rather, the child’s spontaneous understanding of the scene is assessed non-verbally. For example, their gaze has been taken as an indicator of where they expected the agent to search (3, 4, 7). Looking to the correct search location in these non-verbal tasks was shown to correlate with the cortical maturation of a brain region distinct from those involved in verbal ToM reasoning (9). ToM in adults and older children reliably activates the temporoparietal junction (TPJ), the medial prefrontal cortex (mPFC), the precuneus, and the inferior frontal gyri (IFG) (10) and maturation of these regions has been associated with the emergence of verbal ToM reasoning in childhood (9, 11, 12). In contrast, non-verbal action prediction in preschool children correlated with the supramarginal gyrus (SMG) (9), associated with various other social cognitive functions (13–15) and perceptual processes (16, 17). While topologically neighboring the posterior section of the TPJ (i.e., the angular gyrus), involved in ToM, the SMG differs in terms of its functions and brain regions it interacts with. This supports the idea that young children’s non-verbal action prediction relies on different processes than mature ToM and raises the question of which functional role the SMG plays in supporting these early capacities.

Brain regions do not operate in isolation but process information in exchange with other brain regions, and their function may depend on the network they interact with (18, 19). To understand the function of the SMG in young children’s non-verbal action prediction, it is therefore crucial to understand the networks supporting these capacities. During early childhood, brain networks become increasingly interconnected through the maturation of nerve fiber connections between distant network regions, fostering the development of cognitive functions (12, 20–22). For example, increased connectivity of a dorsal nerve fiber connection between two regions involved in ToM (the TPJ and IFG) was found to relate to children’s breakthrough in verbal ToM reasoning at 4 years (11). The nerve fiber pathways supporting younger children’s non-verbal action prediction, however, are not currently known. To reveal the network supporting these capacities, we therefore investigated the maturation of associated fiber connections.

While the regions involved in verbal ToM in older children strongly overlap with the *default network*, the SMG is part of several different brain networks. In particular, it connects to the anterior insula (AI), and together these regions form the core of the *salience network*, involved in attending to salient stimuli (23). Accordingly, the salience network has also been shown to be involved in processing salient social stimuli (24). Based on the different cortical brain regions associated with verbal ToM and non-verbal ToM tasks (9) and their distinct functional networks (19), we hypothesized a dual pathway model underlying the development of our ability to understand others. We predicted that the maturation of a ventral network, with nerve fiber tracts connecting the SMG and AI, as core regions of the salience network, would support correct action prediction in the non-verbal tasks. We expected this ventral network to be distinct from a dorsal network connecting regions involved in mature verbal ToM. Specifically, these include the previously observed dorsal connection between the TPJ and IFG (11) and the cingulum, connecting the precuneus and mPFC as structural backbone of the default network.

To test these hypotheses, we analyzed high-resolution diffusion MRI data from children in the critical period of 3 to 4 years in relation to their behavioral performance in a non-verbal ToM task and standard verbal ToM tasks. We reconstructed anatomical nerve fiber connections and computed indices of their axonal organization and connectivity based on diffusion properties. In particular, as nerve fibers become more myelinated, they become less permeable to water molecule diffusion. As a consequence, mean diffusivity (MD) in the white matter typically decreases, and fractional anisotropy (FA), indexing the directionality of diffusion, typically increases during childhood (25). Further, we used the number of reconstructed streamlines within the fiber tracts as an index of connectivity strength of the tract, which increases during development (26). We then related these micro- and macrostructural measures of connectivity of the fiber tracts with children’s verbal and non-verbal false belief task performance. As ToM development is related to language and executive function development, we additionally assessed children’s linguistic and executive abilities and tested their contribution to the analyzed effects.

## Results

The study and hypotheses were preregistered at OSF (https://doi.org/10.17605/OSF.IO/HMKFD). The analyses follow this preregistration unless explicitly stated otherwise.

### Brain networks for correct action prediction in non-verbal ToM situations

To test whether 3-to 4-year-old children’s performance in non-verbal ToM tasks is indeed associated with nerve fiber tracts connecting regions of the salience network, we reconstructed the hypothesized tract connecting SMG and AI with tractography using the Automated Fiber Quantification Pipeline AFQ (see Methods for details). We then related children’s non-verbal ToM task performance with indices of axonal organization (MD and FA) and connectivity (number of streamlines) in these tracts in general linear models (corrected for multiple comparisons, details see Methods). This revealed a negative association of non-verbal ToM scores with MD in the left SMG-AI connection, in a cluster close to the SMG (N = 40, cluster size = 36, cluster size significance threshold = 26, p = .016 see Figure 1A). In the right hemisphere, the streamline count of the SMG-AI connection showed a positive relation with non-verbal behavioral scores (N = 40, p = .011, Fig. 1B). Both the MD and streamline count effects were independent of children’s language and executive function scores.

**Figure 1.**
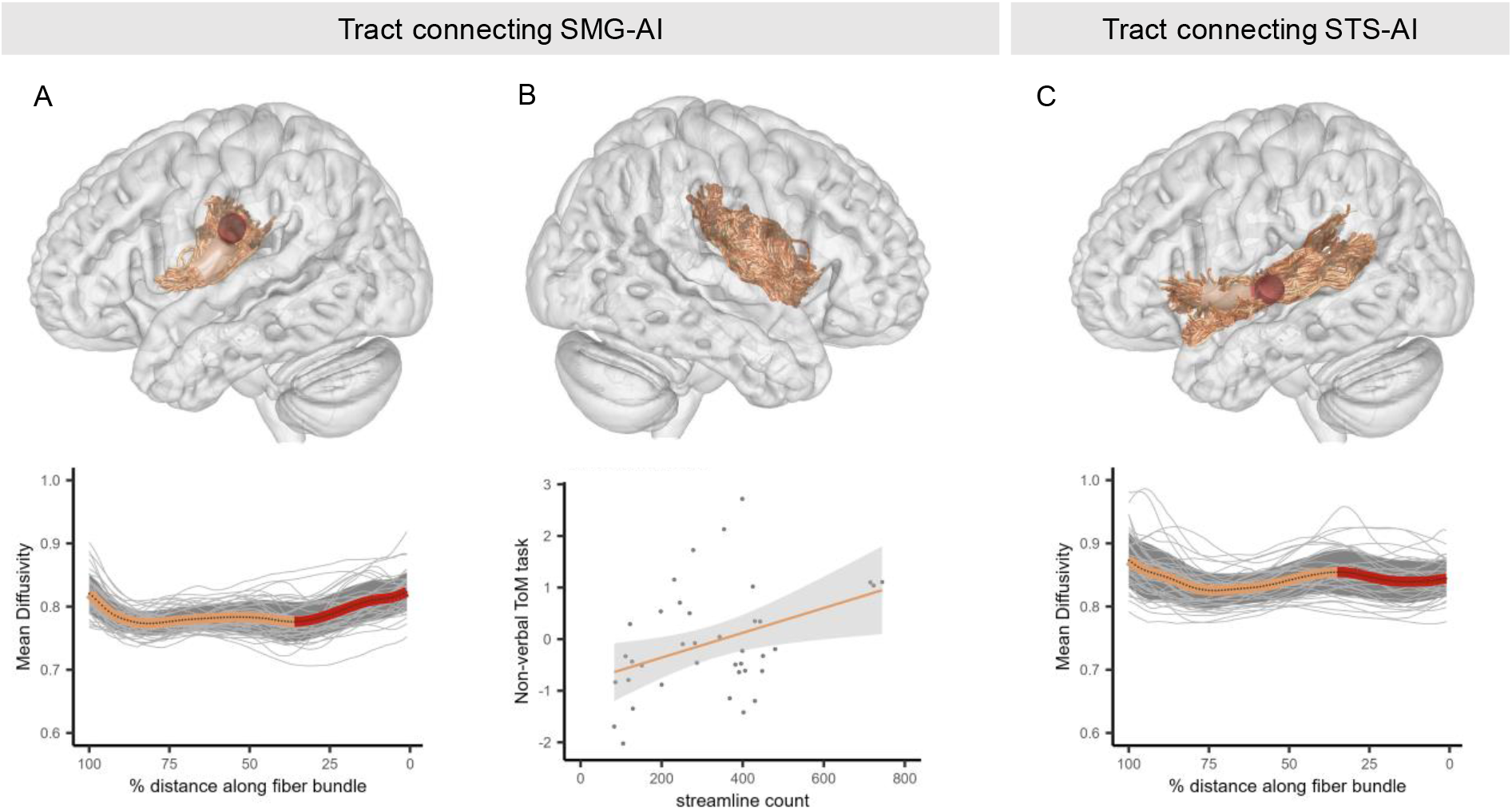
Relations of non-verbal ToM scores and structural connectivity. (A) Significant negative linear relation of MD (in 10^−3^ mm^2^/s) in the dorsal section (red) of the left tract connecting the SMG and AI (orange) with non-verbal action prediction (cluster size = 36, cluster size significance threshold = 26, p = .016). (B) Positive relation of the streamline count of the SMG-AI connection in the right hemisphere with success in the non-verbal ToM task (p = .011). Both SMG-AI effects were independent of language and executive function and of verbal ToM. (C) Significant negative relation of MD (red) in the left STS-AI with non-verbal action prediction (cluster size = 36, cluster size significance threshold = 36, p = .025) in an additional exploratory analysis. The effect was independent of age, language, executive functions, and verbal ToM. All effects along the 100 nodes of each tract (MD and FA) were cluster-size corrected for multiple comparisons with a Bonferroni-corrected significance threshold of p < .05/2. As streamline count yields a single index for the entire tract, these effects had a significance threshold of p < .05. All effects in the top row are visualized on a representative participant. Abbreviations: MD: mean diffusivity, AI: anterior insula, SMG: supramarginal gyrus, STS: superior temporal sulcus.

The SMG and AI are structurally strongly connected to the superior temporal sulcus (STS), involved in processing dynamic social cues, such as gaze (27, 28). Exploratorily, we therefore additionally reconstructed the fiber tract connecting the STS and AI. This showed a significant negative relation of the non-verbal ToM scores with MD in this tract in the left hemisphere (N = 40, cluster size = 36, cluster size significance threshold = 36, p = .025, Fig. 1C). This effect was independent of age, language, and executive function.

### Brain networks associated with verbal ToM

To compare the above results with pathways found for verbal ToM, we re-analyzed children’s verbal ToM data (previously analyzed in (11) using the same analytical approach as for the non-verbal data above. That is, we reconstructed fiber pathways connecting regions of the ToM network, i.e., the arcuate fascicle connecting the IFG and TPJ and the cingulum connecting the mPFC and PC using AFQ. In line with the previous study (11), this analytical approach reproduced a significant relation of verbal ToM with the right arcuate fascicle. Specifically, we observed a negative relation between performance and MD in a temporal cluster of the fascicle (N = 41, cluster size = 21, cluster size significance threshold = 19, p = .008, see Figure 2A). The effect was dependent on the co-developing abilities of language and executive function. Furthermore, we found significant bilateral positive relationships between verbal ToM ability and the streamline count of the cingulum (N = 40, left cingulum: p = .0001, right cingulum: p = .004), which were independent of language and executive function. In addition to these two hypothesized tracts, we reconstructed the short-range connections between the TPJ and PC for exploratory analyses as these are core regions of the ToM network. These showed no relation with verbal ToM performance.

**Figure 2.**
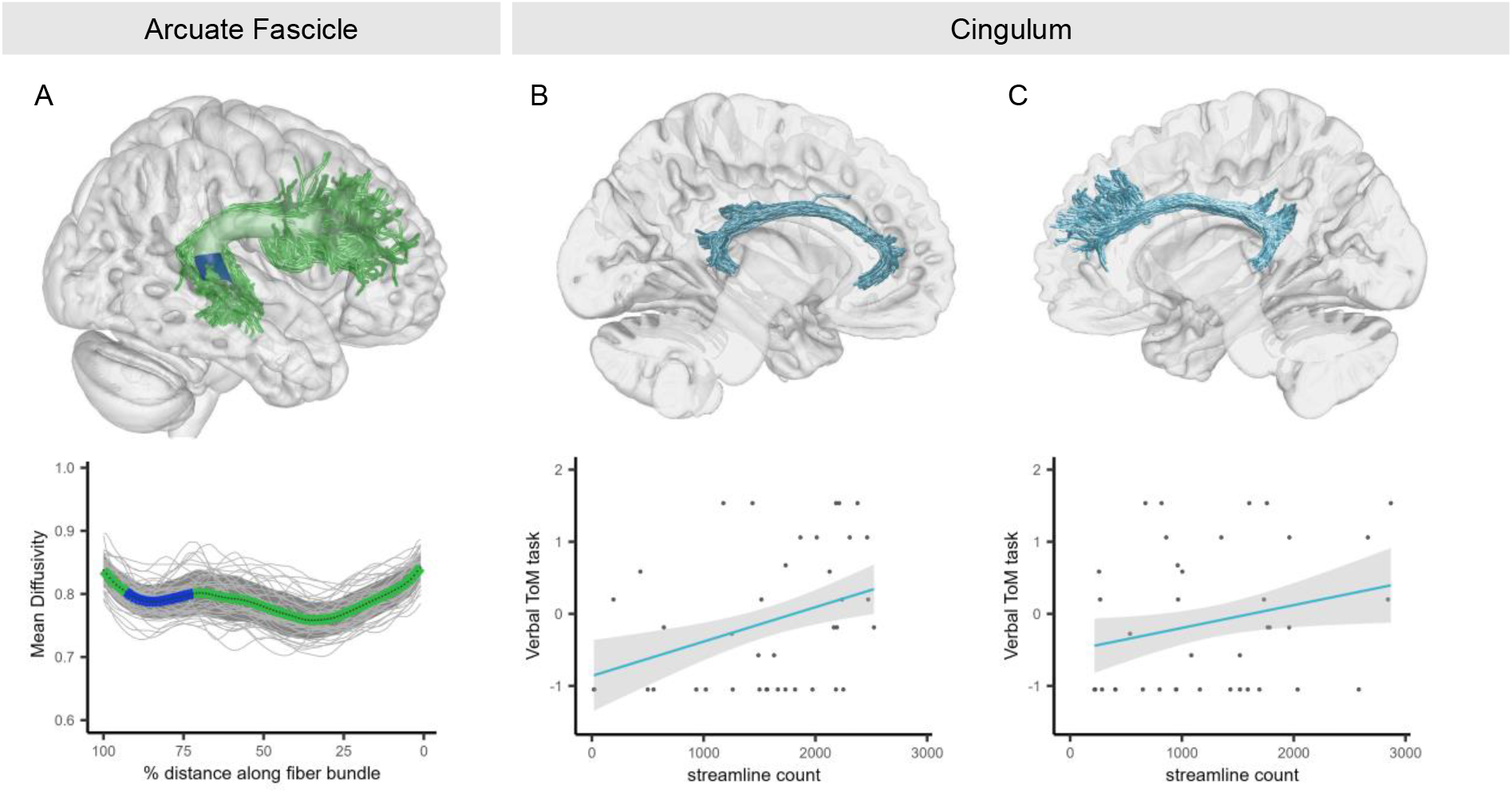
Relations of Verbal ToM scores and Structural Connectivity. (A) Significant negative linear relation of MD (in 10^−3^ mm^2^/s) with verbal ToM in the temporal section (blue) of the right arcuate fascicle (green). (B) Relation of the streamline count of the cingulum (turquoise) in the left (p = .0002) and (C) right (p = .005) hemispheres with verbal ToM. The arcuate fascicle effect was dependent on language and executive function, while the cingulum effects were independent of these factors. All effects were independent of non-verbal action prediction. All effects along the 100 nodes of each tract (MD and FA) were cluster-size corrected for multiple comparisons with a Bonferroni-corrected significance threshold of p < .05/4. As streamline count yields a single index for the entire tract, these effects had a Bonferroni-corrected significance threshold of p < .05/2. All effects in the top row are visualized in a representative participant.

**Figure 3.**
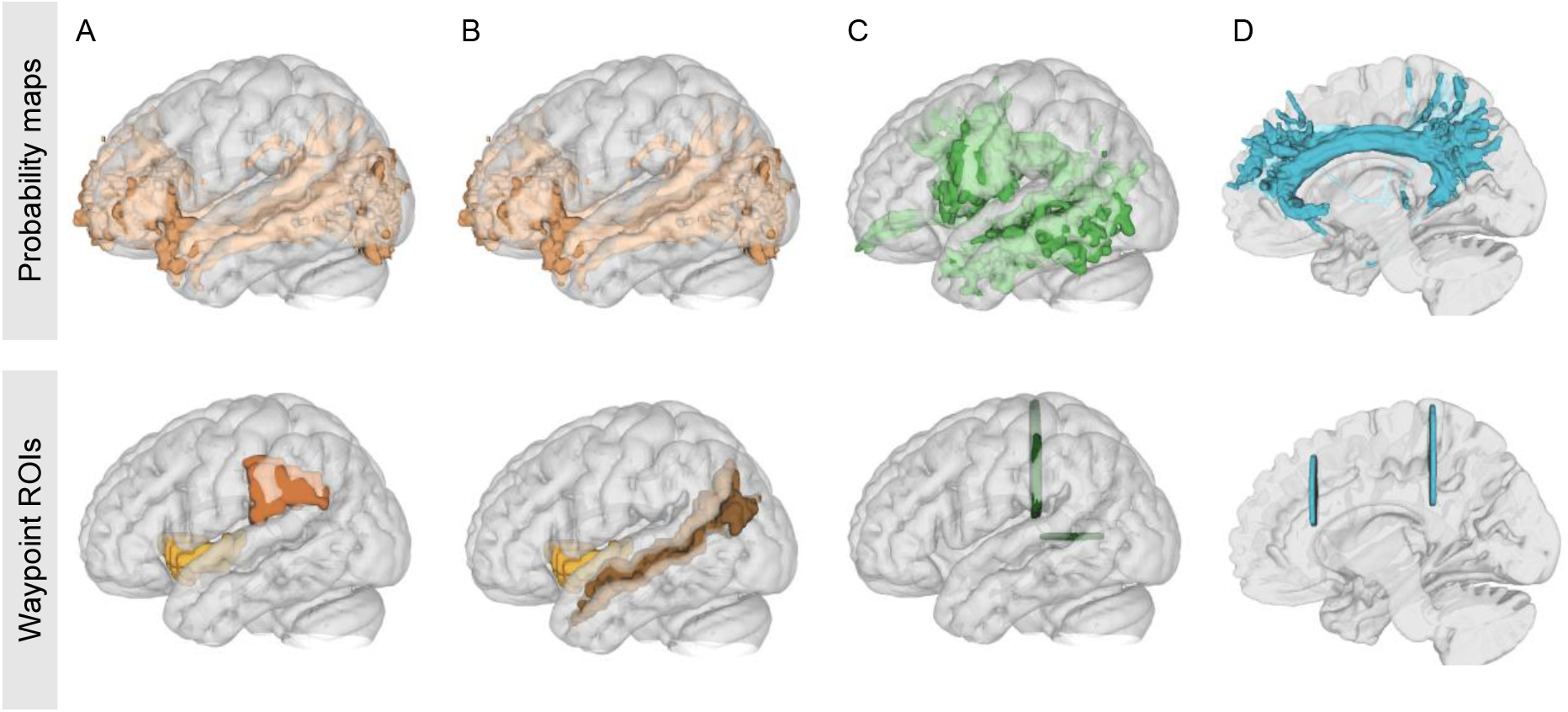
Probability maps and ROIs used in AFQ to define tracts of interest. Probability maps (top row) and waypoint ROIs (bottom row) were used to define tracks of interest. Maps and ROIs are shown for the tracks A) SMG-AI (IFOF map (top), the ROIs for SMG and AI (bottom)), and B) STS-AI (IFOF map (top), the ROIs for STS and AI (bottom)). Furthermore, as implemented in AFQ, the probability maps and waypoint ROIs for the C) arcuate fascicle (green) and the D) cingulum (blue). Visualized in DSI Studio. Abbreviations: AI: anterior insula, SMG: supramarginal gyrus, STS: superior temporal sulcus, PC: precuneus, TPJ: temporoparietal junction.

### Dissociation of verbal and non-verbal ToM tasks

To test whether the effects found for the non-verbal and verbal tasks were independent of one another, we added the respective other behavioral score as a covariate into our linear models. These analyses showed that the effects associated with the non-verbal task in the connection between SMG-AI and STS-I were present independently of children’s verbal ToM performance. onversely, the effects found for verbal ToM in the arcuate fascicle and cingulum were present independently of success in the non-verbal task.

## Discussion

While ToM has traditionally been assumed not to emerge before 4 years, preverbal infants already seem to consider others’ mental states when predicting their actions in non-verbal tasks. This has created a puzzle concerning when ToM develops and what might explain this discrepancy between verbal and non-verbal tasks. We addressed this puzzle by investigating the neural networks associated with young children’s non-verbal action prediction in these tasks and comparing them with the networks supporting the emergence of verbal ToM reasoning between 3 and 4 years. This revealed distinct structural brain networks with a ventral network for young children’s non-verbal action prediction versus a dorsal network for passing verbal ToM tasks. Specifically, non-verbal action prediction was associated with indices of maturation and connectivity of ventral nerve fiber tracts connecting the AI with the SMG and STS, core regions of the ventral attention or salience network (19, 29). In contrast, verbal ToM reasoning was associated with the maturation and connectivity of the arcuate fascicle and the cingulum, two dorsal nerve fiber tracts connecting core regions of the default network (30) and mature adult ToM network.

The nerve fiber connections observed for non-verbal action prediction showed no relation to verbal ToM and vice versa, and the effects were statistically independent of the respective other task type. This confirms that verbal vs. non-verbal capacities rely on distinct and independent networks. These results extend previous findings of a dissociation of verbal and non-verbal capacities in behavior and in local cortical brain structure (9) to the level of brain networks and structural connectivity. Furthermore, they reveal the neural networks and connections associated with non-verbal ToM tasks. A similar distinction of ventral versus dorsal brain networks has been proposed for other domains of cognition, including language (31) and attention (32). In these domains, the ventral network is typically thought to support more perception-based, bottom-up processes compared to higher-order, top-down processes supported by the dorsal network.

Indeed, the salience network, associated with young children’s success in non-verbal ToM tasks, supports bottom-up filtering of salient stimuli that stand out from a stream of sensory input (24). Within this stream of sensory input, social stimuli, such as agents or gaze cues, elicit particularly high saliency (33, 34). Accordingly, the salience network is involved in bottom-up social attention processes, such as detecting social cues and processing others’ emotions or pain (24). Relatedly, its dysfunction has been linked to various social-affective disorders (35). The role of the salience network in the non-verbal tasks suggests that infants’ and young preschoolers’ correct non-verbal action prediction may depend on bottom-up processing of salient social cues, such as the presence and gaze of other agents. Such bottom-up processing would predict that success is context-dependent and influenced by the relative saliency of different aspects of the task, in line with the observed fragility of success in non-verbal ToM tasks (5, 6, 41, 42). Further, this is in line with several recent theoretical accounts suggesting that success in non-verbal ToM tasks may rely on social cues that guide children’s attention and memory (36, 38). Indeed, recent behavioral data show that infants misremember objects in a location cued by the presence or gaze of another person (39), potentially allowing them to predict where the other person will search for these objects in non-verbal ToM tasks. In young children who lack the cognitive capacities for mature ToM, bottom-up social cueing may thus facilitate the understanding of others and ability to learn from them (36, 38).

The salience network for non-verbal ToM tasks were dissociated from the dorsal nerve fiber tracts found for verbal ToM reasoning. Specifically, children’s verbal ToM was linked to maturational indices of the arcuate fascicle (11) and the cingulum. Both tracts connect core regions of the mature ToM network (10), supporting the notion that passing verbal ToM tasks around 4 years indeed constitutes an important developmental step toward mature ToM. The relation of the arcuate fascicle to verbal ToM was dependent on children’s language and executive function. This is in line with the important role of these co-developing abilities in the emergence of verbal ToM (7, 40) and with the arcuate fascicle being a core tract in language processing (31), typically left-lateralized in adults but more bilateral in children (41). Our findings support the cingulum as a structural backbone in the functional ToM network, in line with findings showing that patients with cingulum lesions suffered from deficits in ToM (42). The ToM network strongly overlaps with the default network (43), involved in higher-order social cognitive abilities (44, 45), such as thinking about oneself (46), episodic simulation of the past or future (45), and ToM. As these processes rely on internal representations that are decoupled from direct perceptual input, it has been proposed that the default network sustains internal representations (44). Indeed, such internal representations are needed to flexibly reason about others’ mental representations when these differ from the perceived world, as 4-year-old children begin to do in the verbal ToM tasks.

Our results thus show a dissociation of understanding others in the developing brain. A ventral system is centered around the salience network and relates to non-verbal action prediction for bottom-up, perception-based processes while a dorsal, inference-based ToM system, based on the default network, supports top-down processing of internal, perceptually decoupled representations. A bottom-up, perception-based system for understanding others, as an early alternative to cognitively more effortful ToM, may be adaptive for infants and young children as they are strongly dependent on others and still lacking higher cognitive capacities. This system may not only be useful for young children. It may persist into adulthood where it could provide a fast and efficient way of understanding others, for example, in situations of limited time or cognitive resources (47–49). Indeed, adults also show similar non-verbal behaviors (37, 48, 50, 51) as do non-human primates (52). Future research should therefore investigate whether these behaviors also rely on a ventral salience system rather than the dorsal system recruited for mature verbal ToM.

A further exciting question that arises is how impairments of the salience and default networks relate to individual differences in understanding others, including in clinical conditions with altered social cognitive processing, such as autism spectrum disorder (ASD, 53, 54). Previous research found greater deficits in non-verbal compared to verbal ToM tasks in ASD (55), and differences in attention to social stimuli are a signature of ASD from early in life (56). Future research may therefore want to investigate whether the observed ventral system shows larger alterations in ASD than the mature ToM system (57).

In conclusion, our findings show that non-verbal action prediction and verbal ToM reasoning rely on different neural networks. While the emergence of verbal ToM reasoning in preschool-age children was supported by dorsal fiber pathways connecting core regions of the default network, we found that successfully predicting others’ actions in non-verbal ToM tasks relies on ventral fiber connections between the AI and the SMG and STS, which constitute regions of the salience network. This suggests that, for solving non-verbal ToM tasks, children make use of more stimulus-driven social attention mechanisms, questioning theories of mature representational ToM capacities before the age of 4 years.

## Methods

### Participants

Data from a sample of 3- and 4-year-old children (N = 43), including behavioral assessments and diffusion MRI data, were analyzed for the present study. Two participants had to be excluded due to insufficient reconstruction of white matter tracts and one due to irregularities within all reconstructed tracts, except for the arcuate fascicle. This resulted in a sample of N = 40 participants (N = 41 for the arcuate fascicle) (median 4.08 years; range 3.07 - 4.58 years; 24 female). All children participating in the study had parental informed consent and the study was conducted in accordance with the approval obtained from the Ethics Committee at the Faculty of Medicine of the University of Leipzig.

The present dataset was previously used to address different research questions (9, 11). The question of the fiber pathways involved in non-verbal ToM performance analyzed with AFQ has not been addressed before.

### Behavioral Assessment

Children performed verbal as well as non-verbal ToM tasks. In addition, children’s language and executive function abilities were assessed to test for their potential contribution to effects observed for ToM. This cognitive assessment spanned three separate days, occurring within a median period of 13 days. The tasks were performed in counterbalanced order across participants, except for the verbal ToM tasks, which were always conducted last to prevent any potential influence on the non-verbal ToM task. The tasks are briefly described below; for more details, see (7).

### Non-verbal ToM Task

As a non-verbal ToM task, an anticipatory-looking task was conducted. Here, children were presented with short videos while their gaze was recorded with an eye-tracker. The videos showed a Y-shaped tunnel and an animated agent, e.g., a cat, observing a mouse running through the tunnel and hiding in one of two boxes at the end of each tunnel exit. In false belief trials, the cat left the scene, and – unknown to the cat – the mouse then also left. When the cat came back and entered the tunnel to look for the mouse, the children’s anticipatory gaze at the tunnel exits was recorded as a measure of where they expected the cat to search for the mouse. For the MRI analyses, a non-verbal ToM score was computed as the differential looking score of relative looking times to the correct minus the incorrect box.

### Verbal ToM Tasks

Children participated in two standard false belief tasks, which showed a strong positive correlation (Spearman’s r(43) = 0.879, p < 0.001) and were aggregated to a common verbal ToM score. In the false location task, a candy was hidden in a bag. The child observed the hiding, together with a mouse hand puppet. The mouse puppet then left the scene and the sweet was moved from the bag to a previously empty box. When the mouse returned, the child was asked: (1) where the mouse would search for the sweet, (2) whether the mouse knew where the sweet was, and (3) where the mouse believed it was. A control question ensured that the child remembered the sweet’s actual location. In the false content task, a box of chocolates was presented to the child, and they were asked what they expected to be inside. All kids believed chocolate to be in the box. They were then shown that there were in fact pencils in the box. The mouse puppet showed up, and the children were asked: (1) whether the mouse knew what was in the box, (2) what the mouse believed was inside, and (3) what they had initially believed was in inside, along with a control question about the actual content of the box.

### MRI Data Acquisition & Preprocessing

The dMRI data were acquired using a Siemens 3T TIM Trio scanner. A multiplexed echo planar imaging sequence was used, with an isotropic resolution of 1.9 mm (TR = 4,000 ms, TE = 75.4 ms, b-value = 1,000 smm^-3^, 60 directions, GRAPPA2, 5:32 min). A field map was obtained immediately after the dMRI scan. Additionally, an anatomical scan was acquired using the MP2RAGE sequence with a resolution of 1.2 x 1 x 1 mm (TR = 5,000 ms, TE = 3.24 ms, GRAPPA3, 5:22 min). Before preprocessing the dMRI data, volumes affected by motion artifacts were manually removed. Motion correction was performed by rigidly aligning all volumes to the last volume without diffusion weighting (b0) using flirt58 from the FSL software package. The dMRI data were then rigidly aligned to the anatomical image, which had already been aligned to the MNI standard space and interpolated to 1 mm isotropic voxel space. Distortions were corrected using the corresponding field map. All these transformations were combined and applied to the data in a single step, minimizing the need for multiple interpolations. To prepare the data for AFQ analysis, diffusion data were ACPC aligned.

### Structural Connectivity Analysis with AFQ

To reconstruct white matter fiber tracts from the diffusion data, we used the Python package Automated Fiber Quantification (pyAFQ, 58). Fiber orientation was estimated based on constrained spherical deconvolution and probabilistic tractography was performed with a step size of 0.5 mm and a maximum turning angle of 30°. Then, the desired white matter fiber tracts were segmented with the help of start, end, and waypoint regions of interest (ROIs) and probability maps implemented in AFQ. Table 1 shows an overview of the ROIs used for each studied tract. Exploratorily, we also tried to reconstruct the corpus callosum between bilateral regions of the ToM network in AFQ but this reconstruction did not result in usable tracts, specifically connections between our regions of interest. Therefore, we did not analyze the corpus callosum.

**Table 1.** Region of interest used for reconstruction.

| ToM | Tract | ROIs | Probability maps<br>(implemented in AFQ) |
| --- | --- | --- | --- |
| non-verbal | SMG-AI | SMG (AAL atlas) & AI (Deen Parcellation) | inferior fronto-occipital fasciculus (IFOF) map |
| non-verbal | STS-AI | STS (Destrieux atlas), dorsal ARC ROI used as an exclusion ROI, AI (Deen Parcellation) | IFOF map |
| verbal | Arcuate Fascicle (ARC) | Implemented in AFQ | ARC map |
| verbal | Cingulum (CGC) | Implemented in AFQ | CGC map |
| verbal | PC-TPJ | PC (AAL atlas) & TPJ (Mars Parcellation) | Not used |

Diffusion measures vary along the tract trajectory and are influenced by passing fibers and axons that do not span the entire tract length. Thus, instead of computing average diffusion properties across the entire tract, AFQ calculates the diffusion parameter fractional anisotropy (FA) and mean diffusivity (MD) at 100 equidistant nodes along each tract’s trajectory to localize effects more accurately. In addition to the microstructural measures FA and MD, we computed each tract’s streamline count, which is the total number of streamlines within a tract. All tracts were visually inspected to ensure proper reconstruction. For one subject, all tracts except the arcuate fascicle had to be excluded due to insufficient reconstruction of the tracts with only a few streamlines.

The arcuate fascicle and the cingulum are both included by default in pyAFQ. For custom-defined tracks, we used regions of interest (ROIs) as the start and endpoints in AFQ. Furthermore, for the SMG-AI and STS-AI, we used the IFOF probability map implemented in AFQ to restrict the tracts to ventral connections between these regions. Abbreviations: AI: anterior insula, SMG: supramarginal gyrus, STS: superior temporal sulcus, PC: precuneus, TPJ: temporoparietal junction.

### Statistical Analysis

The obtained measures of structural connectivity from AFQ were correlated with behavioral scores in general linear models with R Studio. Behavioral scores were z-normalized before analysis. To minimize outlier effects, we computed the standard deviations (SD) from the mean and excluded outliers above 3x SD. All effects along the 100 nodes of each tract (MD and FA) were cluster-size corrected for multiple comparisons with a significance threshold of p < .05 and Bonferroni corrected for analyzing two indices MD and FA and the number of tracts investigated. As streamline count yields a single index for the entire tract, these effects had a significance threshold of p < .05 and were Bonferroni-corrected for the number of tracts investigated.

## Acknowledgments and funding sources

This research was funded by a scholarship of the FAZIT Stiftung to CS, and grants by the German Research Foundation (DFG) to PB (project number FR 519/20-2) and to CGW (project number GR 5421/1-2).

## Author contributions

Conceptualization – CS, CGW, and PB. Investigation, Project administration – CGW. Data curation – CS, CGW. Formal analysis, Methodology, and Visualization – CS. Supervision – CGW. Writing, original draft – CS. Writing, review and editing – CGW and PB.

